# Coordination of nuclear RNA processing by speckle-localized kinase TAOK2

**DOI:** 10.1101/2025.09.29.679379

**Authors:** Bridget E Begg, Matthew A Tracey, Shengyan Gao, Svetlana Ernest, Mohd Altaf Najar, George M Burslem, Melanie H Cobb, Beatriz MA Fontoura, Kristen W Lynch

**Author notes:** These authors contributed equally to this work (Bridget E Begg, Matthew A Tracey).

## Abstract

Nuclear speckles are membraneless organelles that act as active splicing hubs especially at sites of high transcription. Emerging views of this dynamic subnuclear structure place it as a hub of RNA processing, impacting steps from transcription to export. To manage this complex microcosm of RNA metabolism, nuclear speckles also require kinases and phosphorylation to execute their functions. The nuclear speckle-localized kinase, TAOK2, mediates the splicing and export of viral transcripts at the nuclear speckle, but its role in the processing of cellular transcripts was unknown. We used siRNA knockdown of TAOK2 and assessed RNA transcripts in both whole cell and nucleocytoplasmic fractions to characterize the complete endogenous effects of TAOK2. We found that TAOK2 knockdown impacts over 10% of the transcriptome, through changes in alternative splicing, export and transcript abundance. Cellular and biochemical phosphoproteomics further revealed nuclear speckle scaffolding proteins SRRM1 and SRRM2 as potential direct phosphorylation targets of TAOK2, mediating its large effects on speckle integrity and speckle-localized splicing. Indeed, knockdown of TAOK2 perturbs almost all speckle-resident serine/arginine (SR)-rich proteins while leaving heterogeneous ribonucleoproteins unperturbed. Altogether, we propose that phosphorylation of SRRM1/2 by TAOK2 plays a structural maintenance role that impacts SR protein-driven exon inclusion at the nuclear speckle.

## Introduction

Nuclear speckles are membraneless, interchromatin condensates first described in 1910 by Ramón y Cajal ((Cajal 1910); summarized in (Lafarga et al. 2009)). For the next hundred years, nuclear speckles were thought to serve primarily as passive storage sites for splicing machinery (Perraud et al. 1979; Spector 1993; Blencowe et al. 1994), but this view of speckles as passive storage sites shifted dramatically in 2012, when Girard et al. (Girard et al. 2012) demonstrated that nuclear speckles house approximately 20% of the nucleus’s catalytically active spliceosomes, indicating that these structures are active hubs of pre-mRNA splicing. Today, much remains to be determined regarding how the structure and activity of nuclear speckles is controlled, and the impact of speckle integrity and function on overall gene expression.

Speckle architecture is scaffolded by the RNA-binding proteins (RBPs) SON and SRRM2, whose intrinsically disordered regions nucleate subdomains of the larger phase-separated body of the speckle (Sharma et al. 2010; Xu et al. 2022). SON and SRRM2 are both members of the larger family of serine/arginine-rich (SR) splicing factors, which as a family are deeply conserved, essential regulators of both constitutive and alternative splicing (reviewed in (Long and Caceres 2009; Shepard and Hertel 2009)) and that are enriched at the nuclear speckle (Caceres et al. 1997). Importantly, speckles are not only sites of splicing, but also impact the regulation and efficiency of the splicing of speckle-resident transcripts. Indeed, disruption of speckles through knockdown of either SON or SRRM2 results in dysregulation of both overlapping and distinct sets of alternative splicing events (Ilik et al. 2020; Zhang et al. 2024), perhaps related to their formation of subdomains within the speckle itself housing different RNA entities. Advances in spatially-resolved RNA sequencing have also revealed that RNAs spliced at speckles tend to be GC-rich with short introns and exons (Barutcu et al. 2022; Tammer et al. 2022; Wu et al. 2024) and frequently possess weaker splice sites or less defined intron boundaries (Wu et al. 2024). Recent quantitative imaging has further demonstrated that pre-mRNAs transcribed from speckle-proximal genes are more efficiently engaged by snRNAs and spliced (Bhat et al. 2024). Examination of transcript dynamics at speckles have also revealed populations of both longer- and transiently-dwelling transcripts at the speckle, suggesting that different transcripts may have different metabolic fates within the speckle.

Although the facilitation of splicing appears to be the predominant function of nuclear speckles, an understanding of the speckle as a site of coordination of multiple gene expression steps is emerging. Speckles are also enriched in proteins related to DNA transcription, replication, integrity, and repair, and it has long been observed that nuclear speckles coalesce at sites of robust transcription, coupling transcription and splicing at highly expressed loci (Carmo-Fonseca et al. 1992; Sacco-Bubulya and Spector 2002; Shopland et al. 2003). However, splicing factors also maintain their interactions at speckles following transcriptional inhibition (Ellis et al. 2008), showing that splicing is facilitated by nuclear speckles independent of the transcription reaction. Moreover, in addition to splicing factors, RBPs that participate in RNA modification, polyadenylation, and nucleocytoplasmic export are also enriched in the speckle (summarized in (Galganski et al. 2017)). It has been shown that intronless genes insensitive to splicing inhibition can be exported via the speckle (Wang et al. 2018), demonstrating that some transcripts transit through the speckle to become competent for nucleocytoplasmic export rather than to undergo splicing. Furthermore, it has been shown that TREX components are required for the release of mRNA from the nuclear speckle (Dias et al. 2010) as a step toward nuclear export.

Kinases and phosphatases that target RBPs are also abundant in speckles and have long been recognized as critical modulators of RNA splicing (Mermoud et al. 1992; Mermoud et al. 1994; Xiao and Manley 1997) reviewed in (Corkery et al. 2015; Naro et al. 2015). Serine/arginine protein kinases (SRPKs) and Cdc2-like kinases (CLKs) phosphorylate members of the SR protein family (Gui et al. 1994; Colwill et al. 1996; Duncan et al. 1998). SR proteins are essential for splicing and their phosphorylation state governs their association with nuclear speckles (Sacco-Bubulya and Spector 2002), incorporation into messenger ribonucleoprotein particles (mRNPs), and subsequent export to the cytoplasm (Huang et al. 2004; Lin et al. 2005). Indeed, overexpression of SRPK1 or CLK1 induces migration of SR proteins away from speckles, speckle dissolution and splicing inhibition (Gui et al. 1994; Colwill et al. 1996; Duncan et al. 1998). Conversely, PP1 phosphatases increase nuclear speckle mRNA retention and cohesion (McIntyre et al. 2025). Together, these findings depict nuclear speckles as dynamically regulated, multifunctional organelles where spatial genome organization, transcriptional regulation, RNA processing, and post-translational control converge.

We recently described a member of another kinase family, TAOK2, a MAP3K-family kinase, as localized to nuclear speckles and required for their structural integrity (Gao et al. 2022). Knockdown of TAOK2 in alveolar lung adenocarcinoma (A549) cells disrupts nuclear speckle structure and causes a concomitant reduction in SON and SC35, essential factors for nuclear speckle organization. In contrast to the activity of SRPKs and CLKs, which dissolve nuclear speckles upon overexpression, TAOK2’s phosphorylation activity promotes the integrity of speckles (Sacco-Bubulya and Spector 2002; Gao et al. 2022). A chemical screen identified TAOK2 as inhibiting the splicing and nucleocytoplasmic export of the influenza virus M transcript, which we determined was post-transcriptionally spliced at the nuclear speckle (Mor et al. 2016; Esparza et al. 2020). A pool of TAOK2 is localized to the nuclear speckle periphery, where it is required for viral M mRNA splicing and export (Gao et al. 2022). The discovery of a kinase whose perturbation impacts both splicing and export provides an opportunity to make inroads into our understanding of phosphorylation-mediated transcript processing at the nuclear speckle.

Here we examine the extent of TAOK2-dependent processing of endogenous RNA and characterize features of transcripts with TAOK2-sensitive splicing and export. We extend our analyses to identify potential phospho-targets mediating TAOK2’s impacts on these processes and synthesize our findings to describe a role for this kinase in the regulation of the SR protein family and the structure of the nuclear speckle.

## Results

### Knockdown of nuclear speckle-localized kinase TAOK2 causes widespread disruption of nuclear RNA processing

Because both the splicing and export of a single viral transcript is dependent on the activity of nuclear speckle-localized TAOK2, we wanted to understand the extent to which TAOK2 activity impacts endogenous RNA processing. To this end, we performed two different RNA-sequencing strategies after siRNA knockdown of TAOK2 (Figure 1A,B). To assay alternative splicing events, we deep-sequenced RNA from TAOK2 siRNA-treated A549 cells and control siRNA-treated cells in duplicate at a depth of ∼60 million reads per sample for robust transcriptome-wide alternative splicing analysis. Alongside, we also used this deep sequencing data to quantify transcripts whose abundances changed after knockdown of TAOK2 compared to siRNA transfection controls to establish the extent to which overall gene expression is changing in response to TAOK2 perturbation. Separately, we also performed nucleocytoplasmic fractionation on siRNA-treated A549 cells followed by RNA-seq on the nuclear and cytoplasmic fractions to quantify transcript abundances before and after TAOK2 perturbation to assess changes in mRNA nuclear export.

**Figure 1.**
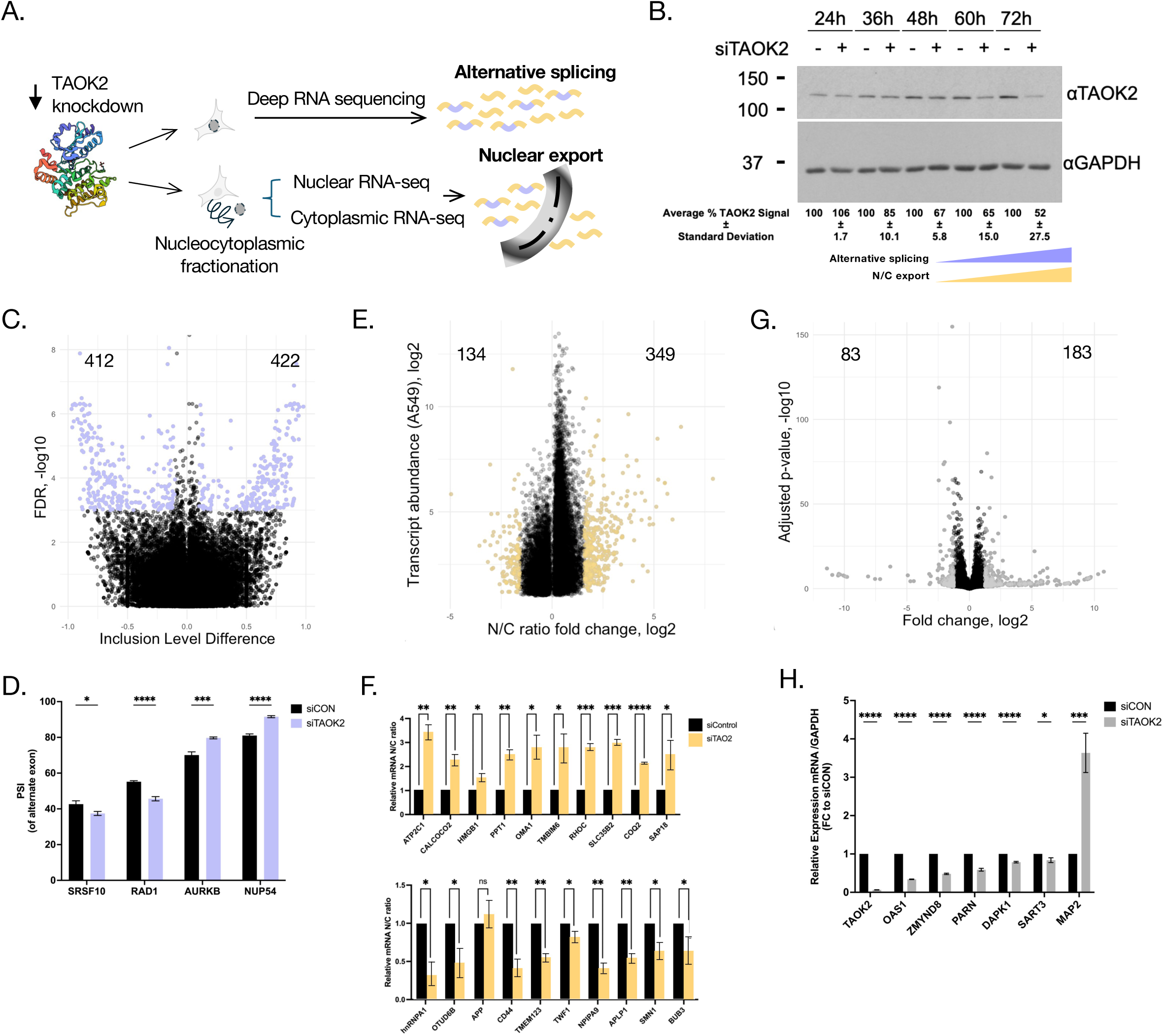
Knockdown of nuclear speckle-localized kinase TAOK2 causes widespread disruption of nuclear RNA processing. A. Experimental workflow for RNA sequencing. B. Western blot of TAOK2 knockdown at 12 hour intervals after siRNA treatment. GAPDH is shown as a loading control. Onset of splicing (purple) and export (yellow) defects after knockdown are indicated with wedges. C. Scatter plot of alternative splicing events upon TAOK2 knockdown. Events with a change in inclusion greater than 15% and an FDR < 0.1 were considered significant and are indicated in purple. D. Validation of key alternative splicing events by ^32^P-labeled RT-PCR at 72h. n ≥ 3. Significance measured by Unpaired t test, * = p<0.05, ** = p<0.01, *** = p<0.001, **** = p<0.0001 E. Scatter plot of changes in nucleocytoplasmic abundance ratios upon TAOK2 knockdown. Transcripts with a greater than 2.5-fold change in abundance were considered significant and are indicated in yellow. F. Validation of top genes whose export is impacted by TAOK2 knockdown by nucleocytoplasmic fractions and qPCR. Relative mRNA N/C ratio upon knockdown (yellow) is normalized to control (black). P-values were calculated using multiple unpaired two tailed Student’s t test. * = p<0.01, ** = p<0.001, *** = p<0.0001 G. Scatter plot of transcript expression changes upon TAOK2 knockdown. Transcripts with a greater than 2.5-fold change in abundance were considered significant and are indicated in grey. H. Validation of key genes with TAOK2-sensitive changes in expression by qPCR at 72h. n = 3. Significance measured by Unpaired t test, * = p<0.05, ** = p<0.01, *** = p<0.001, **** = p<0.0001. A list of total genes and transcripts identified in these analyses are available in **SUPPLEMENTAL TABLE 1**.

We processed our whole-cell deep sequencing data with rMATS (Shen et al. 2014), identifying 2298 cassette exon alternative splicing events occurring in 780 genes with a threshold of >= 20% change in exon inclusion at an FDR <= 0.010 (Figure 1C, purple). We set this conservative threshold for alternative splicing events based on validation with ^32^P-labeled, low-cycle RT-PCR, selecting cutoffs that demonstrated robustness across different genes classes related to RNA processing, cell cycle, and nuclear export (Figure 1D). To determine which transcripts were differentially exported, we manually analyzed our fractionation RNA-seq data. We also sequenced total RNA from TAOK2-depleted cells alongside our nuclear and cytoplasmic fractions to remove from consideration transcripts whose overall expression changes upon TAOK2 knockdown. Following this, we generated per-transcript mean nucleocytoplasmic abundance ratios for knockdown and wildtype cells; 483 transcripts in 451 genes had knockdown N/C ratios that were reproducibly and consistently different from both wildtype replicates at a >= 3-fold change (Figure 1E). We validated by qPCR the top ten transcripts determined to be more retained and more exported and observe results consistent with the RNA-seq (Figure 1F). Finally, 266 transcripts in 243 genes show a >= 3-fold change in abundance upon knockdown of TAOK2, and thus were considered differentially expressed (Figure 1G). Again, we validated these measurements for a subset of individual genes by qPCR with consistent results (Figure 1H). Overall, we find that TAOK2 perturbation results in significant changes in the splicing, expression, or export of >1300 genes (**SUPPLEMENTAL TABLE 1)**, impacting the RNA processing of about 11% percent of the protein-coding transcriptome of A549 cells.

### Genes with TAOK2-sensitive alternative splicing and nuclear export exhibit features of nuclear speckle-associated transcripts

To better understand any connections among RNA processing events impacted by TAOK2, we first asked if the same genes were being impacted at the level of splicing, export, and expression upon knockdown, but find very little overlap among them (Figure 2a). Distinct genes being impacted at the level of splicing and gene expression is consistent with previous observations of RNA-binding proteins, which have suggested that these gene expression steps exist in separate regulatory modules (Martinez et al. 2012; Li et al. 2014; Karlebach et al. 2024). Less expected was the lack of overlap between genes impacted by splicing and export, as we have shown that TAOK2 activity is important for both the splicing and export of an influenza transcript (Gao et al. 2022). This lack of overlap means that upstream splicing changes do not cause the observed transcript export defects. Furthermore, it demonstrates that TAOK2 separately influences splicing, export, and expression. This may occur by the phosphorylation of individual proteins impacting these three processes separately, by a single protein regulating these processes, or by a set of proteins impacting the RNA processing environment more widely.

**Figure 2.**
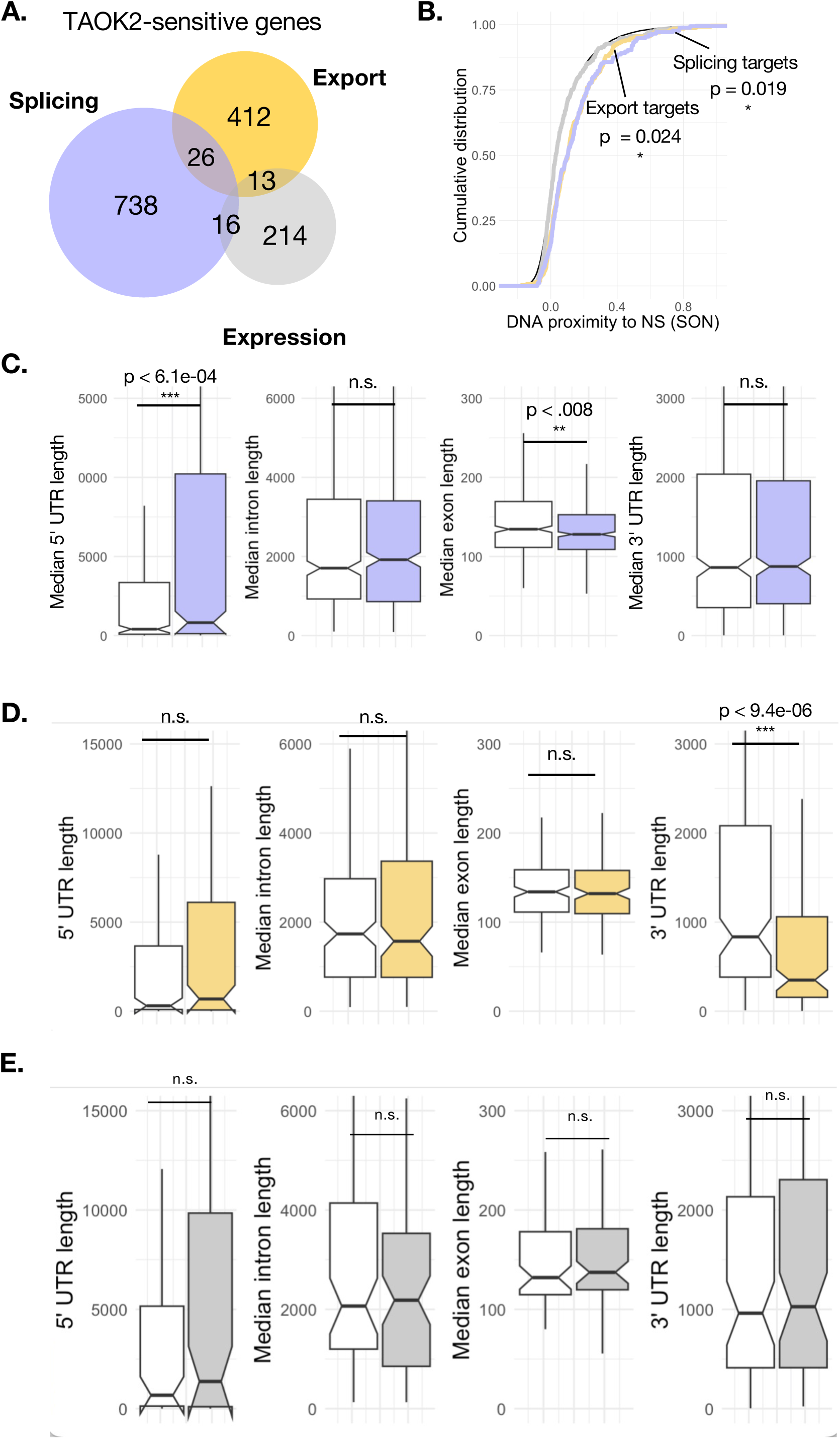
Features of genes with TAOK2-sensitive RNA processing. A. Venn diagram of genes with significant alternative splicing, differential nucleocytoplasmic export, and/or gene expression changes upon TAOK2 knockdown. Purple, alternative splicing; yellow, nucleocytoplasmic export; grey, transcript abundance. B. Distribution of TSA-seq measurements of genes with significant alternative splicing, differential nucleocytoplasmic export, and/or gene expression changes upon TAOK2 knockdown. Purple, alternative splicing; yellow, nucleocytoplasmic export; grey, transcript abundance. Statistical significance was measured by K.S. test. C. Distribution of lengths of 5’ UTR, intronic, exonic, and 3’ UTR gene regions, respectively, of genes with significant splicing events upon TAOK2 knockdown (purple) compared to expression-matched controls (white). Statistical significant was measured by K.S. test. D. Distribution of lengths of 5’ UTR, intronic, exonic, and 3’ UTR gene regions, respectively, of transcripts with significant export defects upon TAOK2 knockdown (yellow) compared to expression-matched controls (white). Statistical significant was measured by K.S. test. E. Distribution of lengths of 5’ UTR, intronic, exonic, and 3’ UTR gene regions, respectively, of transcripts with significant splicing events upon TAOK2 knockdown (grey) compared to expression-matched controls (white). Statistical significant was measured by K.S. test. * = p<0.05, ** = p<0.01, *** = p<0.001, **** = p<0.0001.

It has been shown previously that RNA that is regulated through nuclear speckles also tend to have DNA loci proximal to nuclear speckles, relating gene architecture to RNA processing at the speckle (Barutcu et al. 2022; Wu et al. 2024). We therefore reasoned that genes with RNA processing directly regulated by TAOK2 at the speckle would also have speckle-proximal DNA loci. In fact, we found that genes with TAOK2-dependent transcript export and exon exclusion (and not TAOK2-dependent retention and exon inclusion), have significantly higher SON tyramide signal amplification and sequencing (TSA-seq) scores as compared to expression-matched controls (Alexander et al. 2021), suggesting closer proximity to speckles (Figure 2B). This bias of nuclear speckle proximity for different categories may imply a directionality of TAOK2 or nuclear speckle-influenced activity. This is consistent with TAOK2-regulated splicing and export occurring via the nuclear speckle, as seen for the influenza M transcript (Mor et al. 2016). Conversely, genes with TAOK2-dependent changes in gene expression do not show proximity to the nuclear speckle by this same measurement suggesting that TAOK2’s comparatively limited effects on expression may not occur through speckle-localized processing.

To further probe the connections between TAOK2-regulated RNA processing events, we wanted to know whether the genes impacted at these different steps of RNA processing had similarities at the sequence level, which could support co-residence at the nuclear speckle or suggest other regulatory mechanisms among these groups of genes. We analyzed publicly available gene annotations for genes (in the case of splicing) or transcripts (in the case of export and expression) for well-annotated genes in each group. Genes whose splicing is affected by TAOK2 have longer 5’ UTRs and shorter exons compared to expression-matched controls (Figure 2C). Distinctly, transcripts whose export is impacted by TAOK2 have shorter 3’ UTRs compared to controls (Figure 2D). Shorter introns with higher GC content have repeatedly been shown to be associated with nuclear speckle-localized transcripts (Wu et al. 2024), further connecting TAOK2-sensitive splicing and export with nuclear speckle activity. In contrast, transcripts with TAOK2-responsive gene expression do not show any specific sequence features compared to controls (Figure 2E). Together, these observations strongly suggest that splicing and export RNA processing events are both regulated at the nuclear speckle by TAOK2, though possibly in different regulatory milieus, while genes whose expression levels change in response to TAOK2 knockdown represent a layer of regulation outside of the speckle mRNA regulatory milieu.

### TAOK2’s effects on alternative splicing and nuclear export occur simultaneously

Because the splicing and export of separate sets of endogenous genes appear to be regulated by TAOK2 at the nuclear speckle, we considered two scenarios. First, whether the defective TAOK2-mediated splicing of an export factor caused the TAOK2-sensitive export defects. Second, whether the defective TAOK2-mediated export of a splicing factor caused the TAOK2 splicing defects. We reasoned that if one step of RNA processing drove the other through the defective RNA processing of an intermediate protein, there would be a delay between the effect on one or the other rather than them occurring simultaneously. To answer this question, we performed TAOK2 knockdown over twelve-hour intervals and assessed its effects on some of our validated splicing- or export-defective targets (**SUPPLEMENTAL FIGURE S1**). While we observe initial depletion of TAOK2 protein by 48 hours of siRNA treatment (Figure 1B), TAOK2’s effects on both RNA splicing and export were most apparent at 60-72 hours (**SUPPLEMENTAL FIGURE S1**), when TAOK2 protein is more substantially depleted. Altogether, these results give the impression that the splicing and export impacts of TAOK2 occur simultaneously but are somewhat delayed after TAOK2 knockdown (Figure 1B; splicing (purple), export (yellow)). This supports a model in which one or more proteins phosphorylated by TAOK2 at the nuclear speckle cause simultaneous defects in splicing and export localized to the speckle.

### Phosphoproteomic analysis of nuclear TAOK2 targets

Phospho-targets of TAOK2 have been established in the cytoplasm (Mitsopoulos et al. 2003) (Wojtala et al. 2011) (Nourbakhsh et al. 2021) (Henis et al. 2024), but past work has not investigated nuclear proteins phosphorylated by TAOK2. Since we clearly observe TAOK2 in the nucleus (Gao et al. 2022) and nuclear phospho-targets are most likely to impact splicing and export of mRNAs, we performed nucleocytoplasmic fractionation after siRNA knockdown of TAOK2 and analyzed enriched phospho-peptides with mass spectrometry (Figure 3A). Considering only high-confidence differentially phosphorylated peptides (q-value < 0.01), we found 207 differentially phosphorylated peptides in 142 genes upon TAOK2 knockdown (**SUPPLEMENTAL TABLE 1**). To identify which nuclear processes have phosphorylation most sensitive to TAOK2, we considered the interconnectivity of genes with TAOK2-sensitive phosphorylation using high-confidence, experimentally-determined protein interactions from STRING (Szklarczyk et al. 2023). After removing genes with no interactions among the group, we generated a protein interaction network (Figure 3B). The four largest clusters in the network were associated with RNA splicing (purple), nuclear export (yellow), chromosome organization (blue), and rRNA processing (pink). RNA splicing and nuclear export have established connections to nuclear speckle function, and it has recently been shown chromosome presence at the nuclear speckle periphery can drive and modify overall gene expression (Chen and Belmont 2019; Alexander et al. 2021). rRNA processing, which occurs in another subnuclear organelle, the nucleolus, may also have important transcriptional and nuclear regulatory connections with the nuclear speckle (Shan et al. 2024). Thus, we find that TAOK2 phosphorylation impacts several processes connected to the nuclear speckle, including the critical mRNA processing steps of alternative splicing and nucleocytoplasmic export. Gene ontology analysis of all genes with TAOK2-sensitive phosphorylation also reflects this finding, with significant enrichments for nucleic acid, DNA, chromatin, histone, and RNA binding (Figure 3C).

**Figure 3.**
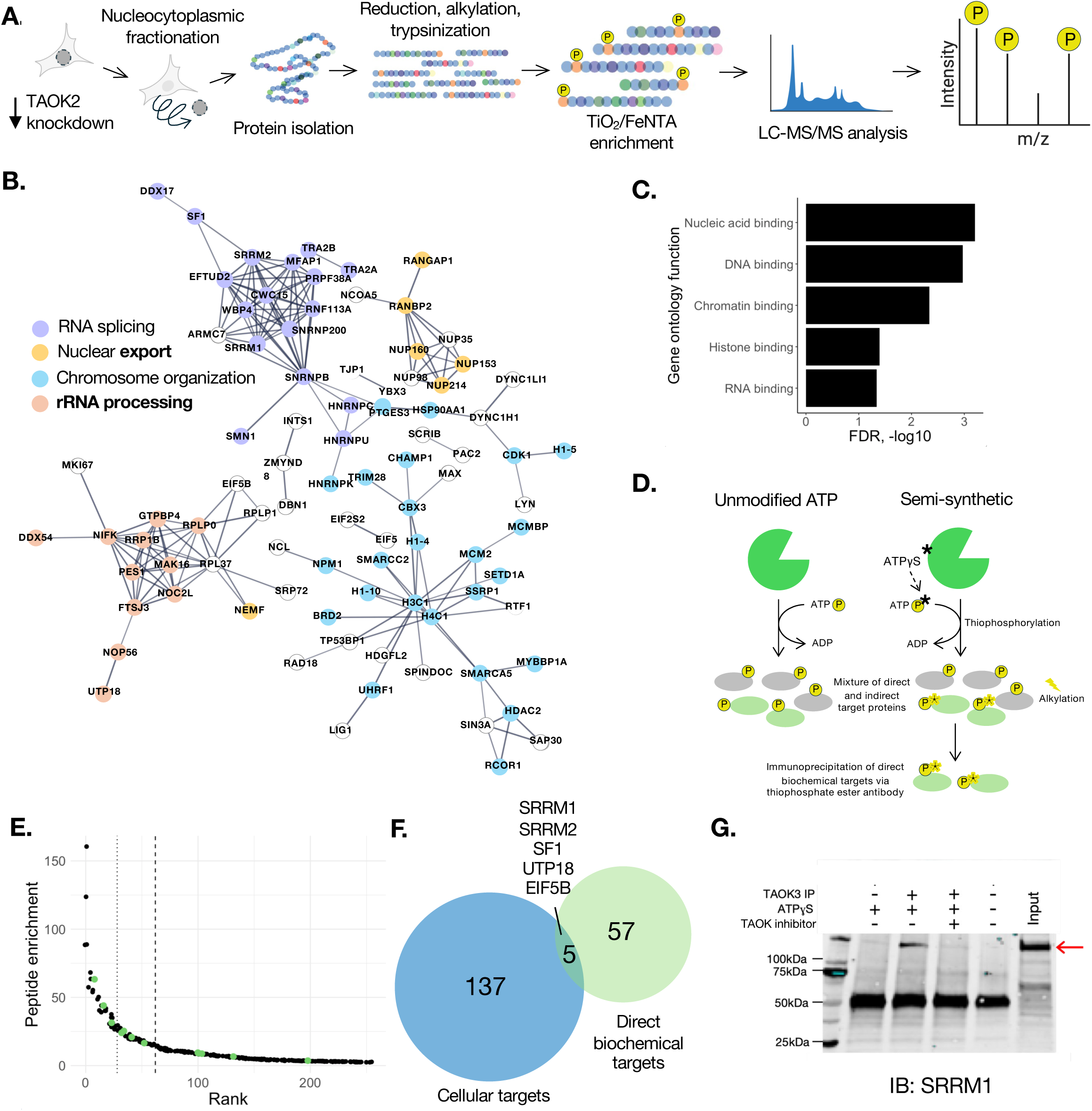
Phosphoproteomic analysis of nuclear TAOK2 targets. A. Experimental workflow for phosphoproteomics. B. Protein interaction network of putative TAOK2 phospho-targets. Protein interactions were reported in STRING in high-confidence experiments or databases. The four most connected modules are indicated by color: RNA splicing (purple), nuclear export (yellow), chromatin organization (blue), and rRNA processing (orange). C. Significant gene ontology functional enrichments of putative TAOK2 phospho-targets. D. Experimental workflow of cell lysate-based analog-sensitive kinase phosphoproteomics. E. Top 250 genes from peptide enrichments derived from TAOK3 analog pulldown and phosphoproteomics. SR proteins are shown in green. The maximum geometric distance is indicated with a dotted line to indicate the elbow of the curve. The top 10% of genes enriched above 1 are indicated by a dashed lines and used for comparison depicted in F. F. Comparison of hits from cellular phosphoproteomic hits from B and top analog-sensitive hits from E. G. Validation of SRRM1. Red arrow indicates SRRM1 protein size.

Because our mass spectrometry from cell nuclei cannot distinguish direct and indirect targets of TAOK2, we also employed a biochemical method to determine potentially direct TAOK2 phospho-targets using affinity purification of thiophosphorylated substrates (Allen et al. 2007). For this approach we used in this case, for technical reasons, the TAOK2 paralog TAOK3, which can employ ATPγS as a substrate and shares >80% identity in the kinase domain with TAOK2. When co-incubated with cell lysate and the ATP analog, the kinase thiophosphorylates its targets, which, after alkylation, can be enriched by immunoprecipitation. Thus, only those proteins modified in the in vitro reactions are identified by immunoprecipitation followed by mass spectrometry (Figure 3D). Proteins phosphorylated in cells will not be recognized using this approach, thereby silencing the background that otherwise makes substrate identification challenging. The top 250 enriched protein targets from this assay appeared rich in SR proteins (green, Figure 3E, **SUPPLEMENTAL TABLE 1**). Ultimately, we considered the top 62 putative biochemical TAOK2 targets in the highest 10% of all protein enriched >1 in the semi-synthetic assay (620 genes) as candidates for TAOK2 phosphorylation (Figure 3F). Comparing these targets with those identified as having impacted phosphorylation in the nucleus upon cellular depletion, we found only five overlapping genes (SRRM1, SRRM2, SF1, UTP18, and EIF5B). Such a limited overlap is not surprising given the different methodologies; however, strikingly, two of the targets identified both as perturbed in cells upon loss of TAOK2 and as direct targets of TAOK in biochemical assays, are SRRM1 and SRRM2 (Figure 3F), key structural components of speckles. While it was unexpected to identify core structural components of the speckle as targets of TAOK2, SRRM1 and SRRM2 are reported to contain multiple phospho-sites highly compatible with TAOK’s substrate specificity (Johnson et al. 2023). We have also further validated SRRM1 as a direct target of TAOK2 by immunoblot (Figure 3G).

### Evidence for regulatory connections among TAOK2 and nuclear speckle scaffold proteins

Because SRRM2 and its binding partner SRRM1 are crucial for the maintenance and organization of the nuclear speckle, we wondered to what extent their expression, and the expression of the other well-established nuclear speckle scaffold protein SON, is impacted by TAOK2 knockdown. Strikingly, by 48h of siRNA treatment, when decreased TAOK2 expression is first evident by Western blot, SRRM1, SRRM2, and SON are substantially downregulated at the protein level by Western blot (Figure 4A). We note that in addition to decreased expression, SRRM1, SRRM2 and SON each also show shifts in migration in the gel that may be indicative of phosphorylation changes (Figure 4A).

**Figure 4.**
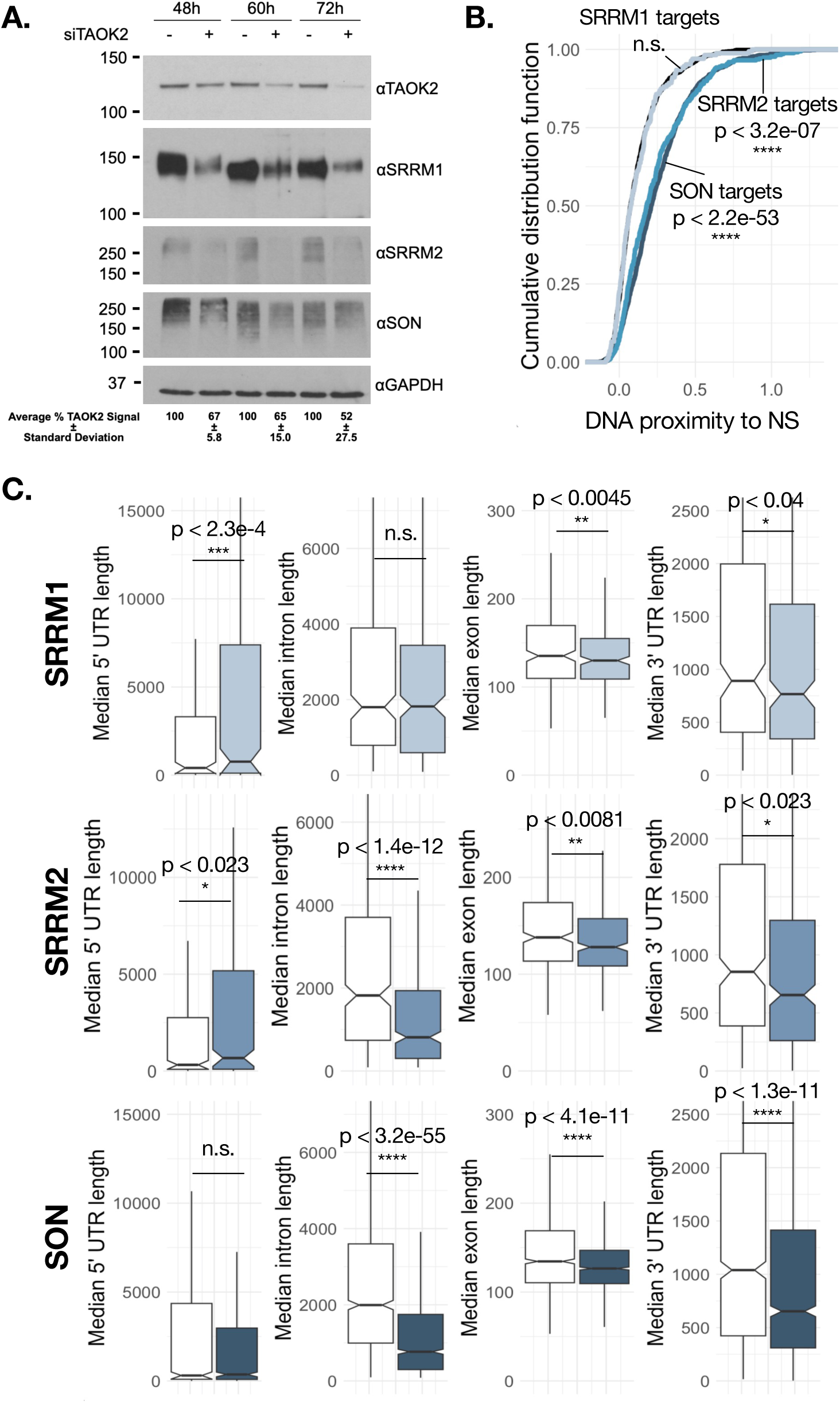
Evidence for regulatory connections among TAOK2 and nuclear speckle scaffold proteins. A. Western blot of nuclear speckle proteins SRRM1, SRRM2, and SON at 48, 60, and 72 hours after TAOK2 knockdown Average fold knockdown and standard deviation of n = 3 TAOK2 knockdown replicates. GAPDH is shown as a loading control. B. Distribution of TSA-seq measurements of genes with significant alternative splicing, impacted by SRRM1 (THP-1), SRRM2 (THP-1), or SON (HEK293) knockdown from available data. Statistical significance was measured by K.S. test. C. Distribution of lengths of 5’ UTR, intronic, exonic, and 3’ UTR gene regions, respectively, of genes with significant export defects upon SRRM1 (light blue), SRRM2 (mid-blue) or SON (dark blue) knockdown compared to expression-matched controls (white). Statistical significance was measured by K.S. test. * = p<0.05, ** = p<0.01, *** = p<0.001, **** = p<0.0001.

To compare the features of genes that exhibit SON- or SRRM2-sensitive splicing with each other as well as with the features of those genes with splicing sensitive to TAOK2, we analyzed data from a recently published study with individual siRNA-mediated knockdowns of SON and SRRM2 in HEK293 cells (Zhang et al. 2024), as well as a published study on SRRM1 and SRRM2 in THP-1 cells (Xu et al. 2022). For both nuclear scaffold proteins SON (dark blue) and SRRM2 (mid blue), but not partner protein SRRM1 (light blue), splicing-sensitive genes had DNA more proximal to the nuclear speckle by SON TSA-seq (Figure 4B), just as for TAOK2. Notably, as with TAOK2, this association is driven by events more included upon knockdown, suggesting a directionality for splicing associated with the speckle. Additionally, all three sets of splicing sensitive genes have shorter median exons and introns, as has been reported for speckle-localized transcripts (Barutcu et al. 2022; Tammer et al. 2022), as well as shorter 3’ UTRs compared to gene expression-matched controls (Figure 4C). Genes with SRRM1-, SRRM2-, and TAOK2-sensitive splicing, however, exhibit substantially longer 5’ UTRs, suggesting that these genes may exist in the same splicing regulatory pathway, somewhat distinct from SON-sensitive genes. Of note, SRRM2 and SON are reported to form separate subphases within the phase-separated nuclear speckle (Zhang et al. 2024), which could explain their processing of different subtypes of genes.

### Impact of TAOK2 knockdown on the expression of core splicing factors and SR/HNRNP proteins

Serine/arginine-rich splicing factors, also known as SR proteins, are essential alternative splicing factors that have long been established as enriched at the nuclear speckle (Caceres et al. 1997). In fact, it has been shown that their phosphorylation patterns are responsive to the assembly and disassembly of nuclear speckles (Gui et al. 1994; Colwill et al. 1996; Duncan et al. 1998; Sacco-Bubulya and Spector 2002). To examine the expression and phosphorylation status of SR proteins more closely, especially those most tightly implicated in speckles or exhibiting overlapping splicing targets with TAOK2, we performed individual immunoblots on SR-family members SRSF1, SRSF3, SRSF5, SRSF6, SRSF9, and SRSF10, and found intriguingly that all exhibited early downregulation and/or shifts in mobility indicative of phosphorylation changes upon TAOK2 knockdown (Figure 5A), with the exception of SRSF9, which shows an increase in expression. We also saw a similar downregulation and/or change in phosphorylation for U170K and U1A proteins of the early spliceosome, at least one of which contains homology to SR proteins (U170K). We also assayed a number of heterogeneous nuclear ribonucleoproteins (hnRNPs), which often function opposite to SR proteins with regard to exon inclusion. Remarkably, we found that hnRNPA1, C, F, L, LL, U, and PTBP1 are, contrastingly, unchanged in abundance or mobility upon TAOK2 knockdown, as were the U2-associated U2AF1 and U2AF2 (Figure 5B). These findings evoke an impact of TAOK2 at the nuclear speckle related to SR-protein regulated exon inclusion of the early spliceosome, possibly by phosphorylation of core scaffolding proteins SRRM1 and SRRM2.

**Figure 5.**
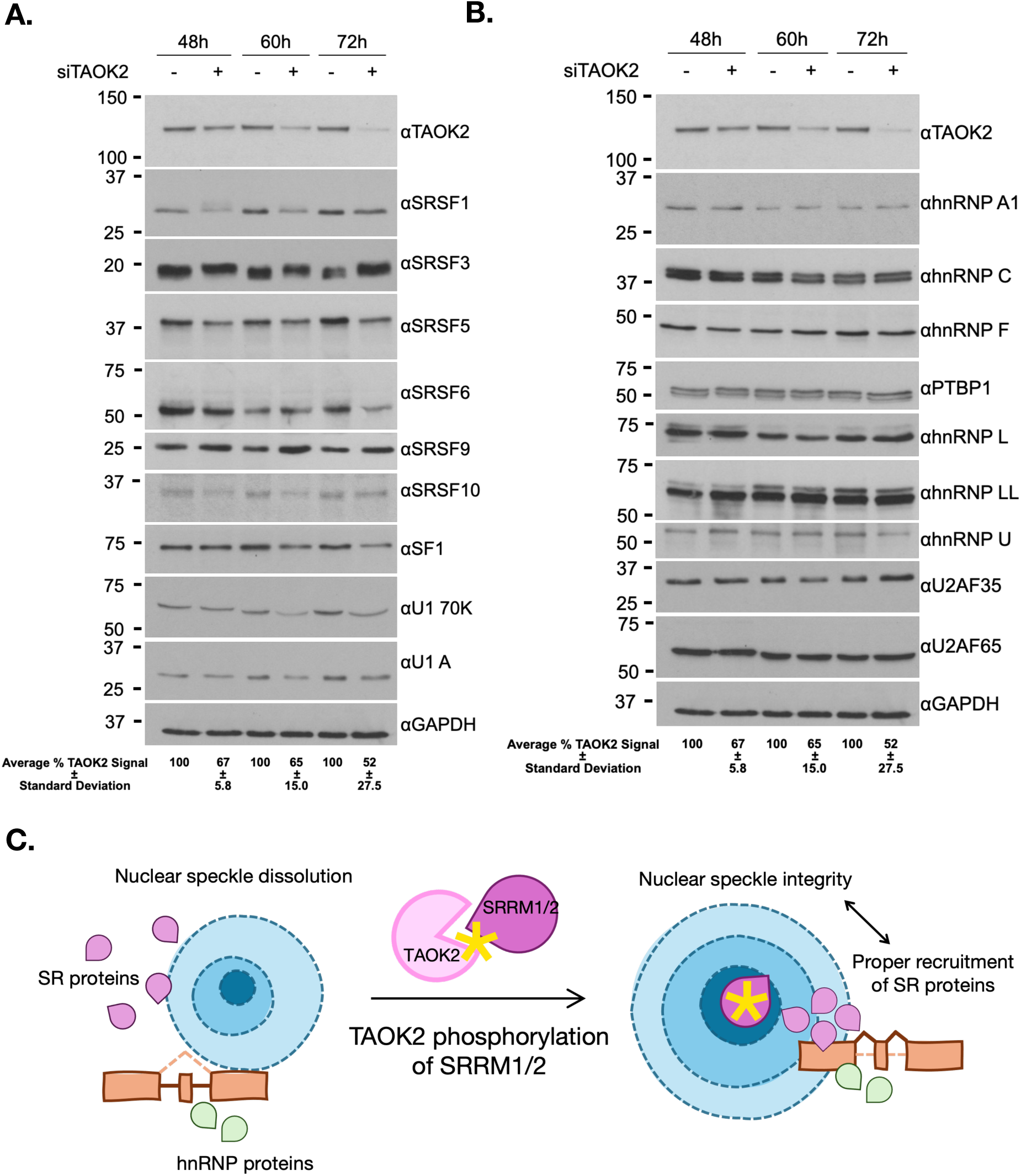
Impact of TAOK2 knockdown on the expression of core splicing factors and SR/HNRNP proteins. A. Western blot of SR and U1 proteins SRSF1, SRSF3, SRSF5, SRSF6, SRSF9, SRSF10, SF1, U170K, and U1A at 48, 60, and 72 hours after TAOK2 knockdown. GAPDH is shown as a loading control. B. Western blot of HNRNP and U2-associated proteins HNRNPA1, HNRNPC, HNRNPF, PTBP1, HNRNPL, HNRNPLL, HNRNPU, U2AF35, and U2AF65 at 48, 60, and 72 hours after TAOK2 knockdown. GAPDH is shown as a loading control. Note for panels A and B that the TAOK2 and GAPDH immunoblots are replicated in Figure 4A for ease of comparison. C. An initial model for the impact of TAOK2 phosphorylation at the nuclear speckle.

## Discussion

Nuclear speckles are currently emerging as active hubs of RNA splicing, but much about their dynamic regulation—and the transcripts that they regulate—still remain to be understood. We show here that TAOK2, a nuclear speckle-localized serine/threonine kinase, is required for the proper mRNA abundance and processing of ∼1000 endogenous genes. Of these genes, >90% are impacted at the level of alternative splicing or nuclear export and these genes are significantly more nuclear-speckle proximal than gene expression-matched controls. In addition, TAOK2-regulated mRNAs share gene features with genes whose splicing and export is sensitive to SRRM1 and SRRM2, known nuclear speckle splicing factors that we observe in two orthogonal phosphoproteomic datasets as putative TAOK2 targets. TAOK2 depletion results in rapid perturbation of not only SRRM1 and SRRM2, but also their partner nuclear scaffold protein SON as well as all classical SR proteins tested, and some U1 factors. Intriguingly, TAOK2 depletion does not appear to impact the expression of hnRNP or U2AF proteins.

While other speckle-associated kinases that phosphorylate classical SR proteins, like CLKs or SRPKs, demonstrate phosphorylation activity that disrupts nuclear speckle formation (Gui et al. 1994; Colwill et al. 1996; Duncan et al. 1998), we have shown that TAOK2 phosphoactivity maintains the structure of nuclear speckles (Gao et al. 2022), and therefore may phosphorylate one or more sites in SRRM1/2 that contribute to nuclear speckle scaffolding or intra-speckle molecular interactions. SRRM2 has recently been shown to contain long, serine-rich charge blocks in its IDR that contribute to intra-speckle interactions with increased phosphorylation (Zhang et al. 2025), a direction of phosphorylation that is compatible with TAOK2 activity. High-throughput reports (Johnson et al. 2023) and our own phosphoproteomics experiments suggest TAOK2 could directly phosphorylate these peptides in SRRM2.

Before their association with speckle scaffolding emerged, SRRM1 and SRRM2 were established splicing coactivators of purine-rich exonic splicing enhancers (Eldridge et al. 1999), alternative splicing sequence motifs known to be regulated predominately by canonical SR proteins and depleted of hnRNPs (Busch and Hertel 2012). It was proposed that U1 components and their associated SR proteins could recruit SRRM1/2, which in turn promoted the recruitment of U2 to facilitate exon inclusion in the alternative splicing reaction (Eldridge et al. 1999). It has also been proposed that SRRM1 alone may act as a general cofactor for canonical SR proteins (Longman et al. 2001), and putative TAOK2-dependent phosphorylation promoting these interactions could have SRRM2-independent effects on nuclear speckle stability in this manner. SRRM1 has also been reported to impact 3’-end processing and unspliced transcript export (McCracken et al. 2002), potentially explaining the mRNA export defects observed upon TAOK2 knockdown. Cytoplasmic TAOK2 has been associated with microtubule attachment and the cytoskeleton (Zihni et al. 2007), so another possibility is that phosphorylation by TAOK2 of SRRM1/2 could create even more upstream speckle architecture by promoting associations with nuclear cytoskeletal proteins.

An intriguing aspect of our work is the sensitivity of SR proteins versus hnRNP protein expression to TAOK2 knockdown. Classically, SR proteins bind splicing enhancer motifs in exons, and interact with U1 and U2 factors to promote exon inclusion. By contrast, hnRNP proteins are typically splicing silencers, blocking interactions with the splicing machinery and resulting in exon skipping (Busch and Hertel 2012). A recent model for splicing at the speckle proposes that SR proteins preferentially inhabit the speckle alongside exonic sequences, while introns and hnRNPs localize just outside the periphery, with spliceosomes situated at the interface at exon/intron boundaries, organizing the splicing reaction (Liao and Regev 2021). Some recent experimental support of this model utilized synthetic reporters with different amounts of RBP motifs to demonstrate that SRSF1 and hnRNPA1 could arrange this way at the nuclear speckle (Paul et al. 2024), but this model has not been proven for other SR and hnRNP proteins or in an endogenous context. Another recent study described even further partitioning of the speckle, with SRRM2 forming a subdomain with U1 proteins and SON forming subdomains with U2 proteins (Zhang et al. 2024).

Our current findings are compatible with these models and meaningfully extend them. We demonstrate that most SR proteins and U1 proteins are sensitive to TAOK2 and nuclear speckle perturbation downstream of TAOK2, while hnRNPs and U2 proteins do not respond to TAOK2— SRRM1/2 speckle perturbation, consistent with a role for hnRNPs outside of the speckle architecture. We also demonstrate sensitivity of U1 and not U2 proteins to TAOK2/SRRM2, which would also support a U2 subdomain separate from U1. Finally, our study may also provide a unifying basis for our understanding of SRRM1/2’s role in the splicing reaction at the speckle. As splicing coactivators enriched in the speckle that can bridge U1 and U2 over introns and SR protein recruitment, SRRM1/2 could orient intronic sequences and hnRNPs to the speckle periphery, and the tightening or loosening of SRRM1/2 interactions of speckle stability would then differentially regulate these relationships, leading to alternative splicing with a bias for exon definition at the short exons enriched at the speckle (proposed model in Figure 5C). Strongly supporting this association is our observation that genes with TAOK2, SRRM2, and SON-sensitive splicing show direction bias in our TSA-seq analysis; not only does this directionality strengthen the association of TAOK2 with these scaffolding proteins, but it is suggestive of a directionality of splicing occurring at the nuclear speckle, which is supportive of some emerging models. While details of this mechanism require further study, we show here that localized kinase activity at the nuclear speckle by TAOK2 promotes core SR proteins to regulate alternative splicing alongside nuclear speckle integrity, and we propose TAOK2 phosphorylation of SRRM1/2 as a potential mechanistic link between these processes.

## Materials and Methods

### Cell culture

Human lung adenocarcinoma epithelial cells (A549) were purchased from ATCC(CRM-CCL-185) and maintained in high-glucose DMEM media (Gibco, 11995-065) supplemented with 10% Fetal Bovine Serum (Gibco, A52567-01) and 100 units/ml Pen/Strep antibiotics (Sigma-Aldrich, A5955) at 37°C with 5% CO_2_.

### RNA interference and transfection

A pool of 3 siRNAs that target TAOK2 (PreDesign siRNA, Sigma-Aldrich) was used for RNA knockdown of TAOK2. Nontargeting siRNAs (a pool of 4 siRNAs, Sigma-Aldrich) were used as control. siRNAs were used at a final concentration of 100 nM. A549 cells were reverse transfected with siRNAs using Lipofectamine RNAiMAX transfection reagent (Invitrogen, 13778150) according to the manufacturer’s instructions

### Cell fractionation and RNA purification quantitative RT-PCR

Cells were harvested by trypsinization and centrifugation. The cell pellets were washed once with cold PBS and pelleted at 300 g for 5 min at 4 °C. The cell pellets were resuspended with 150ml Buffer A (15 mM Tris-HCl pH 8.0, 15 mM NaCl, 60 mM KCl, 1 mM EDTA, 0.5 mM EGTA, 0.5 mM Spermidine, 10 U/mL RNase Inhibitor), incubated for 3 min on ice, then added with 150 ml 2X lysis buffer (0.5% NP-40 added to buffer A), mixed by inversion for 6 times, and incubated on ice for 5 minutes. Cells were then centrifuged at 400 g for 5 min at 4 °C. 200 ml supernatants were taken and stored at -80 °C as cytoplasmic fractions. Pellets were resuspended with 1 ml RLN buffer (50mM Tris-HCl, pH8.0, 140 mM NaCl, 1.5 mM MgCl_2_, 0.5% NP-40, 10 mM EDTA, 10 U/ml RNase Inhibitor) and incubated on ice for 5 minutes, centrifuged at 500 g for 5 minutes and the pellets were served as nuclear fractions. RNA was isolated with RNeasy Plus Mini Kit (Qiagen, 73404).

### RNA-seq of nucleocytoplasmic fractions

RNA from total cell lysates, nuclear or cytoplasmic fractions were firstly run on Agilent Tapestation 4200 to determine the quality and level of degradation. Only RNA samples with RIN score of 8 or higher were used for further analysis. RNA concentration was determined using Qubit fluorimeter prior to starting library preparation. One microgram of DNase-treated RNA was used to prepare library with TruSeq Stranded mRNA Library Prep Kit (Illumina). After library preparation, poly(A) RNA was purified and fragmented for strand specific cDNA synthesis. cDNAs were then A-tailed and ligated with indexed adapters. Samples were then PCR-amplified, purified using AmpureXP beads (Beckman), and validated for quality again using Agilent Tapestation 4200. Samples were then quantified by Qubit and run on Illumina NextSeq 500 using V2.5 reagents for sequencing.

### Calculation of nucleocytoplasmic (N/C) ratios

Transcripts per million (tpms) were estimated using kallisto (Bray et al. 2016) with standard parameters for transcripts annotated in GENCODE Release 49 (GRCh38.p14) for two replicates each of WT or KD nuclear, cytoplasmic, and whole-cell samples for a total of twelve samples. We considered only transcripts expressed > 1tpm in both WT, whole-call samples for estimation of nucleocytoplasmic ratios. To calculate the N/C ratios, we added a pseudocount of 1 to all samples and then calculated initial ratio of the nuclear sample to the cytoplasmic sample per replicate, producing two replicate KD ratios and two replicate WT ratios. Following, we only considered transcripts where either 1) both KD ratios were less than both WT ratios or 2) both KD ratios were more than both WT ratios. We then calculated a mean KD and mean WT N/C ratio for each transcript and considered transcripts with an absolute value log2(fold-change) in N/C ratio of >1.58 differentially exported upon TAOK2 knockdown. Using the whole-cell WT and KD samples, we used DESeq (Love et al. 2014) to assess changes in gene expression upon TAOK2 knockdown and removed any genes with significantly changing expression upon knockdown from the differentially exported set.

### Quantitative RT-PCR of export targets

cDNA was made with iScript cDNA synthesis kit (Bio-Rad, 1708891) from 1.0 mg RNA and 1:5 diluted with nuclease-free water followed by quantitative PCR using LightCycler 480 SYBR green system (Roche) according to manufacturer’s instruction. 18S rRNA was used as an internal control It should be noted that as the calculation of N/C ratios developed, some of the specific validated transcripts dropped out of the final considered list for downstream feature analysis.

### siRNA-mediated TAOK2 knockdown for splicing analysis

An OnTarget Plus SMARTpool (Dharmacon, #SO-3019437G) siRNA targeting TAOK2 (9344) was used for TAOK2 knockdown. A549 cells raised in media were reverse transfected in triplicate with siRNAs using Lipofectamine RNAiMAX transfection reagent (Invitrogen, 13778150) according to the manufacturer’s instructions. siCON control pool #2 (Dharmacon, #D-001206-14-20) was used in parallel for control samples. siRNAs were used at a final concentration of 30 nM. RNA was harvested using Trizol (Invitrogen, #15596026).

### RNA sequencing for splicing analysis

Total RNA extracts assessed by Bioanalyzer with a RIN value > 6.0 and were sent to Azenta for Illumina library preparation with PolyA selection for paired-end, 150bp sequencing at a minimum of 100 million reads per sample.

### Estimation of significantly changing splicing events

Splicing analyses were performed on three replicates each of siRNA-treated TAOK2 knockdown or siCON-treated control A549 cells using rMATSv4.1.2 (Shen et al. 2014) using standard parameters with a STAR genome index built on hg38. Only skipped exon splicing events from genes expressed >1tpm (estimated by kallisto as described) were considered for downstream analyses. Splicing events were further filtered for a significance threshold of FDR < 0.01 and an absolute inclusion level difference of greater than 0.2.

### RT-PCR of splicing targets

Low-cycle reverse transcription (RT)-PCR was performed according to standard methods previously described (Lynch and Weiss 2000, Melton 2007, Agosto 2023). Primer sequences and cycling conditions are available in **SUPPLEMENTAL TABLE 2**. Briefly, RNA was isolated from either control or TAOK2 siRNA treated A549 cells as described above by using TRIzol Reagent (Invitrogen) following the manufacturer’s protocol and was reverse transcribed with Moloney Murine Leukemia Virus Reverse Transcriptase (M-MLV RT) (Promega) using an input of 500 ng of RNA and gene-specific primers. PCR reactions were then performed using ^32^P-labeled primers and were run on 5% denaturing polyacrylamide gels (PAGE). ^32^P-labeled product bands were detected and quantified by densitometry using an Amersham Typhoon Phosphorimager (Cytiva) and Quantity One 1-D analysis software (Bio-Rad). Significance was determined using an unpaired Student t test from at least 3 independent experiments and was performed using Prism 10 software (GraphPad).

### Estimation of gene and transcript expression changes

Gene or transcript expression was estimated in three replicates each of TAOK2 KD and WT samples using kallisto as described. We then used these estimates with DESeq2v1.44.0 in Rv4.4.0. Genes or transcripts with an adjusted p-value of less than 0.05 and an absolute log2(fold-change) of greater than 1.58 were considered significantly differentially expressed.

### qPCR of expression targets

RNA was isolated from either TAOK2 targeting siRNA or control siRNA treated A549 cells as described above and was reverse transcribed with M-MLV RT (Promega), random hexamers (Invitrogen), and an input of 1 μg of RNA. cDNA was diluted 1:10 and used for SYBR-based quantitative (q)PCR (PowerUp SYBR Green Mastermix, Applied Biosystems) in an Applied Biosystems QuantStudio 6 Flex system in a 384-well format. Relative expression of target mRNAs was normalized to GAPDH control and calculated using the ΔΔCT method (Liva 2001). Significance was determined using an unpaired Student t test from 3 independent experiments and was performed using Prism 10 software (GraphPad). Primer sequences are located in **SUPPLEMENTAL TABLE 2**.

### Feature analyses for splicing, export, and expression targets

For each set of genes or transcripts of interest (TAOK2 sensitive splicing, export, or gene expression), a set of gene expression-matched controls was generated in Rv4.4.0 using the MatchIt package (<u>10.18637/jss.v042.i08</u>). Because the directionality of the knockdown effect (i.e. inclusion or exclusion, retention or ectopic export, upregulation or downregulation) sometimes showed different gene expression profiles, an expression-matched control set was established for each set. Except where noted in the text, directionality did not impact the results of the feature analysis. Gene and transcript features were derived from Gencode basic gene annotations from GRCh38 using custom code in Python. Transcripts were quality controlled by checking for the presence of the annotated stop and start codons. We note that this conservative annotation quality control check reduced the overall number of genes considered in each set. Length and GC content features were extracted from GTF files and used for downstream analysis. SON TSA-seq scores were obtained for IMR90 cells from (Alexander et al. 2021). Data for SRRM1 and SRRM1 knockdown was obtained for THP-1 cells in (Xu et al. 2022). Data for SRRM2 knockdown was obtained for HEK293 cells in (Zhang et al. 2024).

### Cell culture and lysis for phosphoproteomics

A549 cells were cultured as described and were treated with phosphatase inhibitors for 1 hour prior to collection via scraping and centrifugation. Cells were resuspended in 10X volume of NIB (15 mM Tris-HCl (pH 7.5), 60 mM KCl, 15 mM NaCl, 5 mM MgCl_2_, 1mM CaCl_2_, 250 mM sucrose) containing protease and phosphatase inhibitors and homogenized by gentle pipetting. Cells were incubated on ice for 5 minutes to lyse outer cell membranes before centrifugation at 1000 rcf for 5 minutes at 4°C. The supernatant contains the cytoplasm and was reserved for further analysis. The pellet was washed by 3 times, by resuspending in NIB buffer containing protease and phosphatase inhibitors and collected by centrifugation at 1000 rcf for 5 minutes at 4°C. Nuclei were further resuspended in 50mM Triethylammonium bicarbonate (TEABC) buffer and sonicated using a probe sonicator at 325 amplitude for total of 12 seconds in three cycles.

### Protein preparation for phosphoproteomics

Prior to processing, the cytosolic fraction was buffer exchanged into 50mM TEABC via Amicon® Ultra 3K device (UFC500396), with 3 cycles of dilution and centrifugation at 12000 RCF for 30 minutes. Protein estimation was performed via BCA assay and an equal amount of protein was taken for further processing (1200 ug). For trypsin digestion samples from both cytosolic and nuclear fractions were first reduced by adding dithiothreitol (DTT) to a final concentration of 5 mM and incubated at 60 °C for 30 minutes. Subsequently, iodoacetamide (IAA) was added to a final concentration of 10 mM followed by incubation for 45 min in the dark at 25 °C. Finally samples were diluted with equal volumes of TEABC. Trypsin (Promega) was added at a 1:50 ratio of protein to trypsin and incubated overnight at 37 °C. Following a digestion check (5 µg of sample underwent C18 clean up were injected into Orbitrap Q-Exactive, results showed >92% of peptides with 0% missed cleavage), the digestion reaction was quenched by slow addition of formic acid until a final concentration of 1% by volume and a pH of 3 is reached.

### Double phosphopeptide enrichment

A double enrichment approach was used to provide optimum phospho-peptide detection. Intial phosphopeptide enrichment was performed using the High Select™ TiO_2_ Phosphopeptide Enrichment Kit (ThermoFisher) following the manufacturer’s instructions. The flow through and washes from the TiO_2_ enrichment were subsequently further enriched using the High Select™ Fe-NTA Phosphopeptide Enrichment Kit (ThermoFisher) following the manufacturer’s instructions. The eluted phosphopeptides from both methods were combined and fractionation with the Pierce High pH Reversed-Phase Peptide Fractionation Kit according to the manufacturer’s protocols. Fractions were dried *in vacuo* and stored prior to analysis.

### LC-MS/MS Analysis

Thermo Scientific Orbitrap Exploris 240 mass spectrometer (ThermoFisher Scientific, Bremen, Germany) connected to the Thermo Scientific UltiMate 3000 HPLC nanoflow liquid chromatography system (ThermoFisher Scientific) was used for data acquisition. Peptide digests were reconstituted in 0.1 % formic acid in water (solvent A) to a final peptide concentration of 500 ng/mL and separated on an analytical column (75 µm × 15 cm) at a flow rate of 300 nL/min using a step gradient of 1–25 % solvent B (0.1 % formic acid in acetonitrile) for the first 100 minutes, 25–30 % for 5 minutes, 30-70 % for 5 minutes, 70–100% for 5 minutes and 1 % for 5 minutes for a total run time of 120 minutes. The mass spectrometer was operated in data-dependent acquisition mode. A survey full scan MS (from m/z 400–1600) was acquired with a resolution of 60,000 Normalized AGC target 300 with an injection time of 50 ms. Data was acquired in topN with 20 dependent scans. Peptides were fragmented using normalized collision energy with 37 % and detected at a mass resolution of 30,000 Normalized AGC target 100 with an injection time of 50 ms. Dynamic exclusion was set for 8 s with a 10 ppm mass window.

### Data analysis

Raw data files were processed using the Proteome Discoverer software suite (version 3.0, Thermo Fisher Scientific) with the SEQUEST HT search engine. Spectral data were searched against the UniProtKB Human reference proteome supplemented with the Common Repository of Adventitious Proteins (cRAP, 125 entries) to account for potential contaminants. Trypsin was specified as the proteolytic enzyme, allowing up to one missed cleavage. Static modifications included carbamidomethylation of cysteine residues (+57.021 Da), while dynamic modifications included methionine oxidation (+15.995 Da), protein N-terminal acetylation (+42.011 Da) and phosphorylation (+79.996 Da) at Ser, Thr and Tyr. Peptide identifications were filtered to a minimum length of seven amino acids. Mass tolerances were set to 10 ppm for precursor ions and 0.02 Da for fragment ions. False discovery rates (FDRs) were calculated using a target-decoy approach with Percolator, and results were filtered to a 1% FDR threshold at the peptide-spectrum match (PSM) level. Data was searched using the LFQ pipeline in Proteome Discoverer to identify and quantify phosphorylated peptides and proteins.

### Network construction and enrichment analysis

Proteins identified in the phosphoproteomics assay as demonstrating a significant reduction in phosphorylation upon TAOK2 knockdown (203 genes, **SUPPLEMENTAL TABLE 1**) were assessed with STRING Database 12.0 (Szklarczyk et al. 2023) to determine the connectivity of the nodes in the network. Only considering the 203 identified proteins, we filtered the interaction network for evidence from only experimental or database evidence at high confidence of the interaction (0.700). Biological process gene ontology terms were used to color the nodes. There was an average node degree of 1.24. Built-in k-means clustering was used to assess gene groups: 1) Nucleosome assembly/epigenetic regulation of gene expression (21), 2) Ribosomal large subunit biogenesis/preribosome (15), 3) Spliceosome/U2 pre-catalytic spliceosome (10), 4) Telomerase holoenzyme complex/HSP90 chaperone cycle for steroid hormone receptors in presence of ligand (10), 5) Nuclear pore complex/Rev-mediated HIV export of RNA (8). Category four includes many DNA and RNA-binding components. We performed a control search using the same parameters for proteins phosphorylated upon TAOK2 KD (236 genes, **SUPPLEMENTAL TABLE 1**) and observed a substantially less interconnected network (average node degree 0.729) and only two clusters with >8 members: 1) Exon-exon junction complex (10) and Ribosome biogenesis (10).

### Gene Ontology analysis

The 203 genes with reduced phosphorylation upon TAOK2 KD were subjected to Gene Ontology Function analysis with GOrilla (Eden et al. 2009). Results were then filtered by FDR < 0.05 Background genes were all genes detected by MS in the nuclear fraction.

### Kinase assay and proteomics

Strep His_6_-TAO3 kinase domain (residues 1-320) was purified from SF9 cells (Chen et al. 1999). Two confluent 10 cm dishes of A549 cells grown in F12K medium supplemented with 10% FBS (ATCC 30-2004) were washed in PBS and lysed on ice with a 27 gauge syringe in 0.25 ml of lysis buffer containing 50 mM Hepes, pH 7.7, 150 mM NaCl, 1 mM EGTA, 10% glycerol, 0.2 mM Na_3_VO_4_, 100 mM NaF, 50 mM β-glycerol phosphate, 10 mM MgCl_2_, 0.1% Triton X-100, 0.2 mM phenylmethylsulfonylfluoride, PhosStop EASYpack (1/10 ml, Roche, #04906837001). The lysate was divided into 4 - 0.15 ml aliquots. Purified TAO3 1-320 (4 μg) was added to 2 of the 4 aliquots. A third aliquot was the kinase-free control and the fourth aliquot was the input control and not further processed. The TAO kinase inhibitor CP-43 (AmBeed #850467-66-2), 5 μM, was added to one of the two aliquots containing TAO3. Kinase reactions were started in the 3 aliquots with 0.33 mM ATPγS (BioLog #A060-05). After 20 min at 30° C, reactions were stopped with 20 mM EDTA on ice. To alkylate the thiophosphates in proteins phosphorylated with ATPγS, p-nitrobenzyl mesylate, 2.5 mM, was added for 2 hr at room temperature with occasional mixing. The samples were then sonicated and 0.35 ml lysis buffer lacking MgCl_2_ was added. Samples were sedimented at 13,000 rpm in a tabletop centrifuge for 10 min. A recombinant rabbit monoclonal antibody (7 μL, Thermo-Fisher, #MA-32345), which recognizes the alkylated thiophosphates, was used to isolate the modified proteins. Protein A-agarose beads (Pierce, Protein A/G Plus # 20423) were used to recover the antibody-bound modified proteins and were washed twice with 0.6 ml MgCl_2_-free lysis buffer. Proteins were released with 40 μL 5X SDS electrophoresis sample buffer and loaded onto a BioRad 10% polyacrylamide gel (# 4561033) and electrophoresed until material penetrated 0.5 cm into the separating gel. After staining with Coomassie blue, the gel was sliced and the band was submitted to the UT Southwestern mass spectrometry core for protein analysis.

### Validation of phosphotargets

Aliquots of the four samples, three of which were processed by mass spectrometry, were resolved on BioRad 4-15% polyacrylamide gradient gels (#4568083), transferred to nitrocellulose and immunoblotted with a polyclonal rabbit antibody to SRRM1 (Proteintech #12822-1-AP ).

### Western blot/Immunoblot and quantification

Whole cell protein extract (WCE) was generated by lysing A549 cells in radioimmunoprecipitation assay (RIPA) buffer (0.15 M NaCl, 1% NP-40, 0.5% sodium deoxycholate, 0.1% SDS, 50 mM Tris pH 8) following either control or TAOK2 siRNA treatment as described above at the indicated time point. Protein levels were assessed by loading 5-20 μg of WCE into 8%-12.5% 37.5:1 bis-acrylamide SDS PAGE gels depending on target protein size, with the exception of SRRM1. SRRM1 protein levels were assessed by loading 10 μg of WCE into 3%-8% gradient Tris-Acetate gels (Invitrogen). Proteins separated on SDS PAGE gels were transferred to PVDF membrane (Sigma Aldrich) using Bjerrum Schafer-Nielsen transfer buffer with SDS (48 mM Tris, 39 mM glycine, 20% methanol, 0.0375% SDS) either at 4°C overnight in a tank transfer system (30 V constant, 16h, Bio-Rad) or at room temperature using a semidry blotting apparatus (Fisherbrand) following the manufacturer’s protocol depending on target protein size. Proteins separated on 3%-8% gradient Tris-Acetate gels (Invitrogen) were transferred to PVDF membrane (Sigma Aldrich) using a Bicine and Bis-Tris transfer buffer with SDS (25 mM Bicine, 25 mM Bis-Tris, 1 mM EDTA, 10% methanol, 0.0375% SDS) at 4°C overnight in a tank transfer system (30 V constant, 16h, Bio-Rad). Antibodies used for Western blot were diluted in 5% BSA-TBST and can be found in **SUPPLEMENTAL TABLE 2**. Relative TAOK2 protein levels following 3 independent siRNA knockdown experiments were determined by densitometry using Quantity One 1-D analysis software (Bio-Rad) and normalized to a GAPDH loading control.

## Acknowledgements

We thank members of the Fontoura, Cobb, and Lynch Labs for interesting discussion. This work was funded by NIH R01 AI125524 to KWL, BMAF, and MHC and NIH R35 GM142505 to GMB. MAT was also supported by grant T32GM007229.

## Author contributions

BEB performed experiments, analyzed data, and wrote the manuscript. MAT performed experiments and analyzed data. SE, MAN, and SG performed experiments. MHC, GMB, and BMAF supervised the study. KWL supervised the study and wrote the manuscript.

