## Supplemental Figure 1 for "Coordination of nuclear RNA processing by speckle-localized kinase TAOK2"

**Supplemental Figure 1.** A. A549 cells transfected with siRNAs targeting TAOK2 or non-targeting siRNA control. Cells were collected at indicated time points after transfection. mRNAs from total cell lysates were subjectedto [32]P-labeled quantitative RT-PCR and indicated mRNAs were detected with specific primers. Graphs are mean +/- SEM of three independent experiments. P values were calculated using multiple unpaired two tailed Student's t test. * p < 0.05, ** p < 0.01, *** p < 0.001, **** p < 0.0001.

**A.**

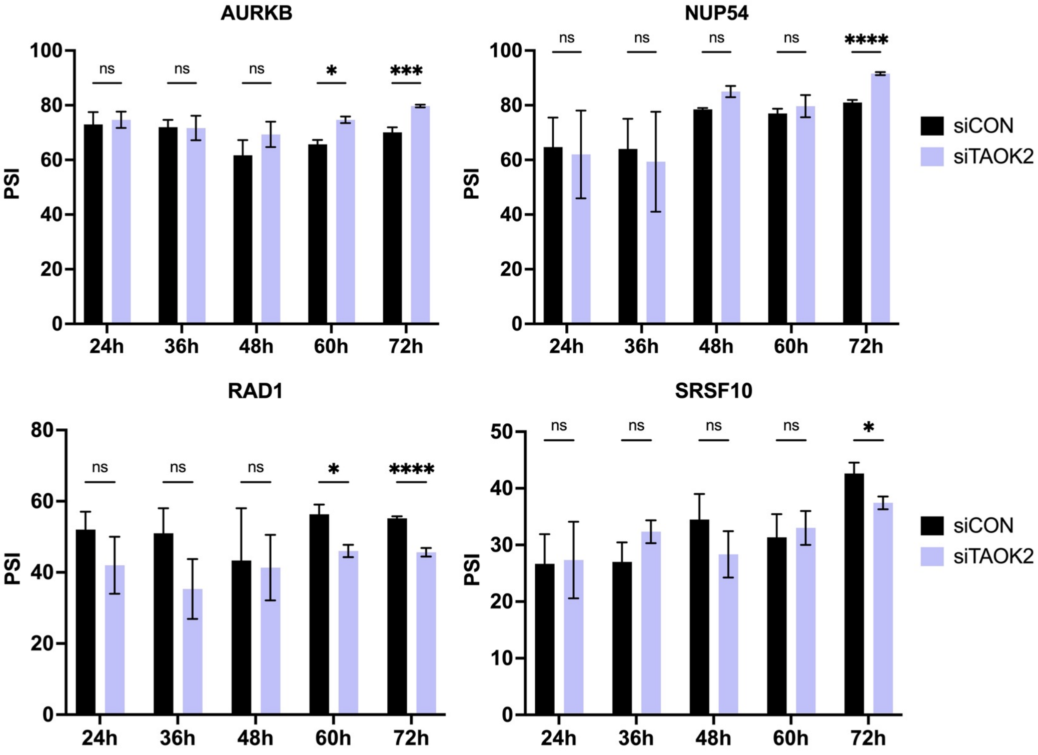


B. A549 cells transfected with siRNAs targeting TAOK2 or non-targeting siRNA control. Cells were collected at indicated time points after transfection and subjected to nuclear-cytoplasmic fractionation. mRNAs from total cell lysates, cytoplasmic and nuclear fractions were subjected quantitative RT-PCR and indicated mRNAs were detected with specific primers. Relative nuclear/cytoplasmic (N/C) ratio of each mRNA was normalized to siRNA control. Graphs are mean +/- SD of three independent experiments. P values were calculated using multiple unpaired two tailed Student's t test. * p < 0.05, ** p < 0.01, *** p < 0.001, **** p < 0.0001.

**B.**

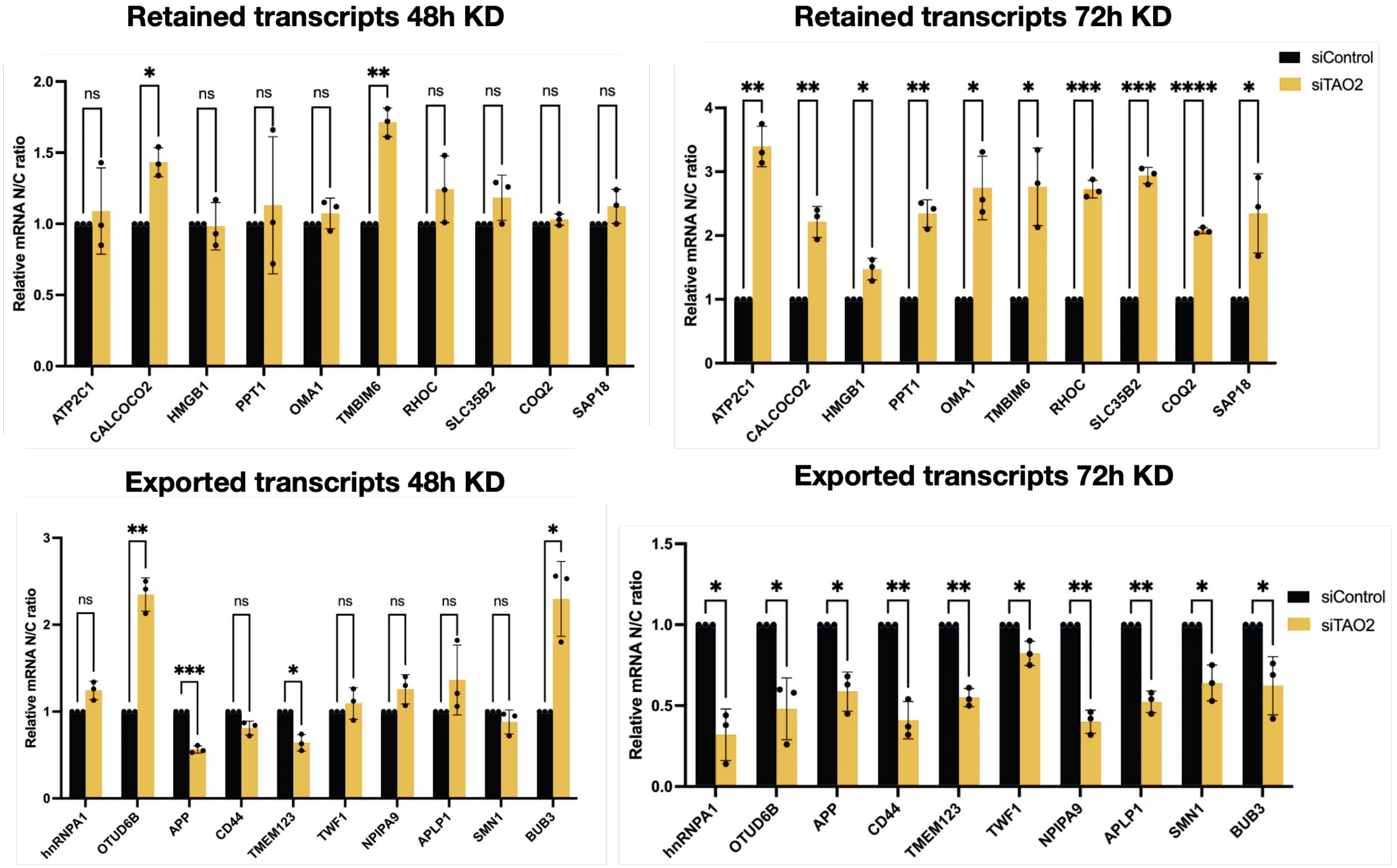
